# Neural Phoneme Processing in Children with and without Dyslexia

**DOI:** 10.1101/2025.10.13.682108

**Authors:** Marlies Gillis, Jill Kries, Jan Wouters, Laura Gwilliams, Maaike Vandermosten

**Affiliations:** Experimental Oto-Rhino-Laryngology, Department of Neurosciences, KU Leuven, Belgium; Laboratory of Functional Anatomy, Université libre de Bruxelles, Brussels, Belgium; Laboratoire de Neuroanatomie et Neuroimagerie translationnelles, UNI – ULB Neuroscience Institute, Université libre de Bruxelles, Brussels, Belgium; Department of Psychology, Stanford University, Stanford, CA, USA

**Keywords:** Dyslexia, Phonological Processing, Neural Tracking, Temporal Generalization

## Abstract

This study investigates the neural dynamics of phoneme processing in 7-year-old children with and without dyslexia (25;9 ♂), using EEG recordings collected during continuous speech listening. By applying temporal generalization to phonetic descriptor decoding, we can disentangle whether potential phoneme processing deficits are due to the maintenance of phonemes in verbal short-term memory and/or inferred differences in phonetic processing speed, both of which are thought to be impaired in dyslexia. We investigated whether phonetic processing depends on the phoneme’s position or its lexical competition.

Our results reveal two key findings that may help explain the challenges faced by children with dyslexia. First, these children exhibit reduced decoding accuracy for word-onset phonemes, suggesting disruptions in either predictive, word-level anticipatory mechanisms or in the intrinsic rhythmic processing aligned with word boundaries. Second, they exhibit increased decoding accuracy for non-onset phonemes with low lexical competition approximately 400 ms after phoneme onset. This pattern suggests that children with dyslexia retain linguistically less relevant sounds longer in verbal short-term memory and process them more slowly compared to their typical reading peers.

Together, these findings suggest that dyslexia is characterized by altered phonetic encoding strategies, specifically inefficient prioritization of relevant phonological information. This work provides new insight into the neural mechanisms underlying phonological deficits and contributes to a deeper understanding of the cognitive basis of dyslexia.

**Significance statement:** Dyslexia is associated with difficulties in phonological processing. Investigating EEG during continuous speech listening, we show that children with dyslexia exhibit weaker encoding of word-onset phonemes and prolonged processing of less informative phonemes. These altered encoding strategies suggest inefficient prioritization of linguistic information, offering new insight into the neural basis of dyslexia.

## 1 Introduction

Dyslexia is a specific neurodevelopmental disorder affecting about 7% of the world’s population (Peterson and Pennington, 2012). Individuals with dyslexia experience difficulties in reading and writing that cannot be explained by factors such as low IQ, poor schooling, or lack of motivation. These persistent struggles often lead to poorer educational and occupational outcomes, along with reduced self-esteem and well-being (Livingston et al., 2018).

Although dyslexia manifests as difficulty with written language (reading and writing), the underlying issue is thought to stem from reduced sensitivity to features of *spoken* language, namely phonemes, the smallest units of speech (Snowling, 2000). The origin of this phoneme processing deficit is speculative; possible hypotheses are poor temporal and auditory processing (e.g. Hämäläinen et al., 2012; Poelmans et al., 2012; De Vos et al., 2017), degraded phonological representations (Hulme and Snowling, 1992; Swan and Goswami, 1997) or issues with accessing or maintaining phonemes in verbal short-term memory (Ramus and Szenkovits, 2008; Boets, 2014).

Behavioural and neuro-imaging studies interested in understanding this link have primarily tested how simple sound fragments, such as isolated phonemes, syllables or non-speech sounds, are processed (for behavioural studies, see review Noordenbos and Serniclaes, 2015, and references within) (for fMRI studies, see Boets et al., 2013; Vandermosten et al., 2020) (for ERP studies, see review Volkmer and Schulte-Koerne, 2018, and references within). However, such simple and artificial stimuli do not fully capture the complexity of phoneme processing in natural, continuous speech, where neighboring phonemes must be processed rapidly and efficiently. On average, a phoneme lasts approximately 100 ms, whereas the neural processing of a phoneme in natural speech can extend up to 400 ms (Gwilliams et al., 2022; Brodbeck et al., 2022; Gillis et al., 2021). Consequently, the brain must process multiple neighboring phonemes in parallel during natural speech perception (Gwilliams et al., 2022; Brodbeck et al., 2022). This parallel and rapid processing of phonemes is believed to be particularly difficult for individuals with dyslexia, given that it requires verbal short-term memory and fast sound processing, two processes which are both reported to be impaired in persons with dyslexia and believed to be at the basis of poor phonological representations (Ramus and Szenkovits, 2008). Therefore, testing whether the neural dynamics of phoneme processing in continuous speech are affected in dyslexia is an important next step.

Phoneme processing of continuous speech can be investigated by means of *neural tracking* (for reviews, see Brodbeck and Simon, 2020; Gillis et al., 2022), the phenomenon where the brain rhythmically follows or tracks features of continuous, natural speech. The extent of neural tracking can be assessed using a variety of methods, ranging from simple linear models, which approximate the relationship between the stimulus and the measured neural responses with an oversimplified linear assumption, to complex artificial neural networks, which can capture non-linear interactions but make the interpretation of these relationships more challenging (Puffay et al., 2023). Most commonly, studies assess neural tracking to investigate the acoustic processing of speech, using acoustic speech features such as the speech envelope or spectrogram. However, phonemes are also robustly tracked by the brain (such as Gwilliams et al., 2022; Gillis et al., 2021; Brodbeck et al., 2022).

Studies investigating neural tracking of the speech’ acoustics in persons with dyslexia generally draw the same conclusion: adults and children with dyslexia track the speech envelope less strongly than their peers (e.g. Di Liberto et al., 2018; Destoky et al., 2020, 2022; Klimovich-Gray et al., 2023), and similar reductions are observed in methods investigating the neural synchronization between the neural responses and speech envelope (e.g. Molinaro et al., 2016; Mandke et al., 2022). Yet, most work has focused exclusively on the speech envelope (except Di Liberto et al., 2018; Klimovich-Gray et al., 2023), which does not directly capture the dynamic processes during phonemic processing. Investigating phoneme processing, Di Liberto et al. (2018) reported atypical neural tracking in the right hemisphere of children with dyslexia, suggesting altered phoneme processing compared to both age- and reading-level-matched peers. Yet, Di Liberto et al. (2018) did not provide any insights into the temporal aspects of phonemic processing.

However, investigating the temporal aspects of phonemic processing may provide valuable insights into the neural dynamics of phoneme processing and how these differ between individuals with and without dyslexia. Such knowledge can clarify the role of verbal short-term memory and the speed of phoneme processing in accessing and maintaining the phonological representations. Another temporally structured aspect worth investigating is whether phoneme processing depends on its position in the word, i.e., whether word-onset phonemes are processed differently from non-onset phonemes. By comparing word-onset and non-onset phoneme processing, one can reveal differences in both top-down and bottom-up phoneme processing mechanisms. Word-onset phonemes play a crucial role in word recognition because they strongly constrain the set of activated lexical candidates. This lexical activation, in turn, is a key mechanism in generating predictions about upcoming sensory input, aligning with predictive coding frameworks. The extent to which dyslexia impacts predictive coding processes remains unclear and underexplored. However, some suggest that individuals with dyslexia might rely on higher predictive coding mechanisms (Stanovich, 1980; Cavalli et al., 2016), serving as a protective factor to compensate for less accessible or degraded phoneme representations. Hence, highly predictable phonemes may rely less on bottom-up auditory processing, often impaired in dyslexia, and more on predictive coding. Following this reasoning, phonemes might be processed differently based on their lexical relevance, i.e., whether they are more or less predictable. In contrast, studies employing amplitude-modulated sound fragments at the word rate (*∼*4 Hz) to examine bottom-up acoustic processing have shown that individuals with dyslexia exhibit reduced neural synchronization compared to typically developing peers (e.g., Hämäläinen et al., 2012; Poelmans et al., 2012; De Vos et al., 2017). This reduction suggests difficulties in tracking the intrinsic rhythms of speech, particularly at the word level. Thus, investigating the temporal dynamics of neural phoneme processing may provide critical insights into the mechanisms underlying phonological difficulties in dyslexia.

Our study uniquely addresses this question by employing a temporal generalization approach. This method provides insight into when and how phonemes are represented in the brain, allowing inferences about the stability of neural phonetic representations. Furthermore, it enables quantification of the duration over which a phoneme is processed, reflecting its maintenance in verbal short-term memory. It also allows for quantifying the rate at which phonetic information propagates across neural populations, thereby indicating the speed of phoneme processing. Using this temporal generalization approach, Gwilliams et al. (2022) demonstrated that during continuous speech, the brain encodes multiple phonemes in parallel and that neural phonemic representations dynamically evolve over time. Since children with dyslexia are thought to have less accessible or degraded phoneme representations (Ramus and Szenkovits, 2008), it is particularly important to determine whether and how differences in phoneme processing duration and speed contribute to their difficulties. The temporal generalization approach presents a promising avenue for addressing this question.

Using the temporal generalization approach, the present study investigates the neural dynamics of phoneme processing in children with and without dyslexia to gain insight into how phoneme representations are maintained in the brain and the speed of phoneme processing. However, this approach has been validated so far only in adult neural data. Specifically, Gwilliams et al. (2022) applied it to 2 hours of magnetoencephalography (MEG) recordings while participants listened to continuous speech. More recently, Kries et al. (2024) demonstrated that the approach is also feasible with electroencephalography (EEG) data, despite its lower signal-to-noise ratio compared to MEG, and with a substantially reduced recording time (25 minutes) in older adults. The next crucial step is to validate this method in children’s EEG data, which is inherently noisier than that of adults, and to use a reduced recording time since collecting more than one hour of EEG data in children is not feasible. If successful, this would provide an objective, neural-based way to assess phonological processing, offering several advantages over traditional behavioral tests. In particular, children can remain engaged with a naturalistic story, thereby reducing attention demands. The absence of explicit task instructions also minimizes fatigue and avoids the ceiling and floor effects often observed in behavioral assessments.

Our study has three main aims:

1. To test whether phoneme-level neural dynamics can be measured in children using EEG over relatively short recordings (17 minutes).

2. To examine whether there are differences in phoneme processing between children with and without dyslexia and whether these are related to their position or lexical competition.

3. To explore whether the differences in phoneme processing are linked to individual variation in reading skills.

## 2 Methods

### 2.1 Participants

Our sample consists of 25 children tested at the beginning of 2nd grade (mean age: 87 *±* 3 months; ∼7 years, 3 months; 16 ♀, 9 ♂). All were native Dutch speakers with normal hearing, no reported neurological or developmental disorders (including autism or attention disorders), no history of brain injury, and no ongoing therapy for speech or language problems. All children participated in a longitudinal intervention study (for more information, see supplementary material, Section S.1). The children were originally included in kindergarten because they were identified as having an elevated cognitive risk for dyslexia. Risk was defined as scoring below the 30th percentile on at least two of three reading-related skills (phonological awareness, letter knowledge, rapid automatized naming). In addition, children scoring below the 30th percentile on phonological awareness and rapid naming also had to score below the 40th percentile on letter knowledge. All 25 children in the current study were in the passive control group and did not receive intervention. For details on the screening procedure, see Van Herck et al. (2023) and Verwimp et al. (2020).

At the beginning of 2nd grade, children completed several behavioral tasks. EEG recordings took place during the summer holiday before the start of 2nd grade (which starts in September). Importantly, the COVID-19 pandemic began in early 2020, with school-closures when the children were in the third trimester of 1st grade. In Belgium, where the data were collected, school closures had a substantial impact on reading and spelling development (Duroisin et al., 2021). However, in a similar longitudinal cohort, the phonological delay seen at the start of second grade, likely due to school closures, had disappeared by the start of third grade (Blockmans et al., 2025).

Reading outcomes were assessed at the start of 3rd grade, when children could be classified as either typical readers or as having dyslexia. Classification was based on standardized word reading (Brus and Voeten, 1973; Verhoeven, 1992), pseudoword reading (Van den Bos et al., 1994), and spelling tasks (Deloof, 2006). Children were classified as dyslexic if they scored below the 10th percentile on at least one task and below the 25th percentile on all other tasks. Thus, dyslexia classification relied on 3rd-grade data, while the reading-related behavioral measures (see Section 2.2) used in this study were collected in 2nd grade, concurrent with the EEG experiment. This approach was chosen because reading measures in the 3rd grade provide a more robust diagnostic marker. Moreover, given the disruptions from COVID-related school closures, 3rd-grade outcomes are likely more reliable than 2nd-grade data for determining whether a child was dyslexic (Duroisin et al., 2021; Blockmans et al., 2025).

Of the 25 children, 9 were identified as having dyslexia (7♀, 2 ♂; mean age 86 *±* 4 months). For group comparisons, we created an age-matched control group of 9 typical readers (5 ♀, 4 ♂; mean age 87 *±* 3 months). This ensured equal group sizes and minimized age-related biases.

The study was approved by the Medical Ethical Committee of the University Hospital of Leuven, KU Leuven, and written informed consent was obtained for all participants (Katholieke Universiteit Leuven; approval number B322201836276).

### 2.2 Reading-related behavioral measures

Among the behavioral tasks completed by the children, we selected four with particular relevance as precursors of reading development. These tasks measure auditory processing, phonological awareness, and phonological verbal short-term memory; skills linked to dyslexia.

#### 2.2.1 Auditory processing

Lower prereading auditory processing skills, particularly those involving low-rate dynamic information, are associated with weaker phonological and literacy development later on (Vanvooren et al., 2017). These skills involve identifying temporally varying cues, which are critical for processing the speech envelope.

To assess this, we used a **rise time discrimination task**, which measures sensitivity to differences in amplitude modulation, specifically the rise time (the duration from sound onset to peak amplitude). In each trial, children heard three sounds: two identical reference stimuli and one deviant with a different rise time. They indicated which sound was different. By adaptively decreasing the rise time difference, we determined the threshold at which the child could still reliably discriminate the deviant.

Stimuli were 800 ms long, created from speech-weighted noise with a fixed linear fall time of 75 ms and variable rise times (15 to 699 ms in 50 steps). The shortest rise time (15 ms) was always the reference and was presented twice per trial. Stimuli were presented unilaterally to the right ear at 70 dB SPL in a three-alternative forced-choice oddity paradigm using the APEX software (Francart et al., 2008). An adaptive staircase procedure estimated the threshold corresponding to 70.7% correct responses (Levitt, 1971). Each child completed the task three times (one practice, two test runs), and the final threshold was taken as the lowest of the two test runs. For details, see Van Herck et al. (2023).

Children and adults with dyslexia generally show reduced sensitivity to amplitude modulations, i.e., higher detection threshold of rise time differences, a finding consistently replicated (e.g. Hämäläinen et al., 2005; Poelmans et al., 2011; Vanvooren et al., 2017). Moreover, detection ability appears to vary continuously with reading skill: better readers are more sensitive to amplitude modulations (Goswami et al., 2002). The distribution of thresholds in our sample is shown in Figure 1.

**Figure 1.**
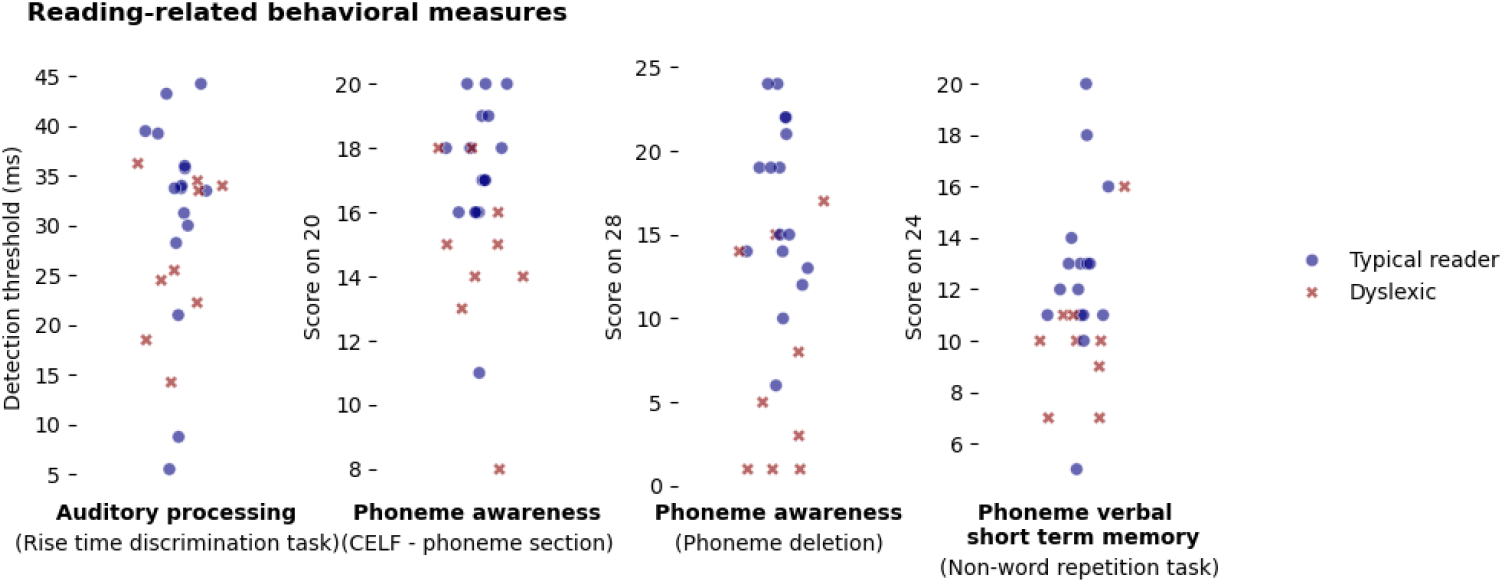
Distribution of the scores on the reading-related behavioral measures of all 25 children, typical reading (annotated by the blue dot) and dyslexic (annotated by the red cross).

#### 2.2.2 Phoneme awareness

Phoneme awareness is the ability to recognize and manipulate phonemes in spoken words, a skill essential for reading and spelling. Dyslexia is strongly associated with deficits in this domain (Snowling, 2000). We measured phoneme awareness using two tasks.

First, a **subtest of the Clinical Evaluation of Language Fundamentals (CELF-4NL)** (Kort et al., 2010), which assesses knowledge of Dutch sound structures through 20 items involving identification, segmentation, blending, and deletion of phonemes.

Second, a **phoneme deletion task**. In each trial, children heard a pseudoword and a target phoneme, and were asked to repeat the pseudoword without the target phoneme. The task included three blocks of eight pseudowords (24 trials in total), with each block focusing on a different type of phoneme deletion and preceded by two practice trials (for details, see Poelmans et al., 2011).

Both tasks reliably reveal lower phoneme awareness in children and adults with dyslexia (Poelmans et al., 2011). Score distributions for both measures in our current sample are shown in Figure 1.

#### 2.2.3 Phonological verbal short-term memory

Phonological verbal short-term memory refers to the temporary storage and manipulation of phonemes, a capacity critical for speech comprehension and reading acquisition (Gathercole and Baddeley, 1993).

We measured this ability with a **nonword repetition task** (Gathercole et al., 1994), using a shortened, Flemish adaptation of the Dutch version (Scheltinga, 2003). The task contained 24 nonwords of increasing length. Children were instructed to repeat each nonword as accurately as possible (for more details on this shortened version, see Vanden Bempt et al., 2021).

Children and adults with dyslexia typically perform worse on nonword repetition tasks than typical readers (reviewed in Melby-Lervåg and Lervåg, 2012). Successful repetition requires analyzing and segmenting the nonword into phonological units stored in long-term memory. Thus, difficulties may reflect impaired access to these representations (Snowling, 2000). Interestingly, a meta-analysis by Melby-Lervåg and Lervåg (2012) found only a weak-to-moderate relationship between nonword repetition and phoneme awareness, suggesting that nonword repetition engages both phonological and nonphonological processes. Score distributions in our current sample are shown in Figure 1.

### 2.3 Experimental protocol: Speech stimulus & EEG acquisition

Children listened to a Dutch story, *De Wilde Zwanen*, narrated by a male speaker. The story lasted approximately 21.5 minutes and was divided into six parts. The first four parts (4 to 5 minutes each) were presented in silence, while the final two shorter parts (about 2 minutes each) were presented with background noise. Stimuli were delivered bilaterally through ER-3A insert phones (Etymotic Research Inc., IL, USA) using the APEX software platform (Francart et al., 2008) at a fixed intensity of 60 dB SPL.

During the experiment, children sat in a soundproof room within a Faraday cage. EEG signals were recorded with a BioSemi ActiveTwo system (BioSemi, Amsterdam, NL) at a sampling rate of 8192 Hz. A fixation cross was shown on a screen throughout the listening task to minimize eye movements.

For the present study, only the first four parts of the story were analyzed, since background noise can alter neural response patterns. Thus, the analyzed neural data consisted of approximately 17.5 minutes of continuous speech presented in silence.

### 2.4 Phoneme Classification

Phoneme-level information was extracted in three steps: alignment, classification according to pronunciation, and classification according to lexical competition.

#### 2.4.1 Phoneme alignment

First, the onsets of the phonemes were determined using the web-based tool of Kisler et al. (2017). By linking the audio files with the input transcripts, the tool generates a phoneme alignment per audio file. More specifically, we utilized its pipeline without automated speech recognition (pipeline specifications: *G2P* → *MAUS* → *PHO2SYLL*), setting the language to Dutch (NL) and enabling the use of a forced aligner. This produced a phoneme alignment, i.e., a TextGrid file with phoneme-level time stamps aligned to the speech signal.

#### 2.4.2 Phoneme classification by pronunciation class

Second, each phoneme was described according to five pronunciation classes proposed by King and Taylor (2000):

- **Phonation**: presence of vocal cord vibration. Descriptors: *voiced* (e.g., /b/) vs. *unvoiced* (e.g., /p/)
- **Roundness**: lip rounding during articulation. Descriptors: *rounded* (e.g., /w/) vs. *unrounded* (e.g., /f/)
- **Manner of articulation**: airflow pattern through the articulators. Descriptors: *long vowel, short vowel, fricative* (e.g., /s/), *nasal* (e.g., /n/), *occlusive* (e.g., /p/), *approximant* (e.g., /r/)
- **Place of articulation**: articulator location. Vowels: *low* (e.g., /a/), *mid* (e.g., /e/), *high* (e.g., /i/) Consonants: *coronal* (e.g., /l/), *dental* (e.g., /f/), *labial* (e.g., /b/), *velar* (e.g., /r/), *glottal* (e.g., /h/)
- **Tongue front–backness**: tongue position. Descriptors: *front* (e.g., /m/) vs. *back* (e.g., /k/)

Phoneme descriptors that had an occurrence count smaller than 5% of the total amount of phonemes were not considered. These descriptors are all related to place of articulation, i.e., *low* for vowels, *dental* and *glottal* for consonants.

#### 2.4.3 Phoneme classification by lexical competition

Third, phonemes were classified according to their role in lexical competition, quantified by cohort entropy. Cohort entropy reflects the uncertainty among candidate words consistent with the phoneme sequence heard so far. High entropy indicates strong competition (many possible words remain), while low entropy indicates weak competition (few words remain). Thus, phonemes with higher entropy are linguistically more informative.

Cohort entropy was computed as the Shannon entropy of the active cohort:

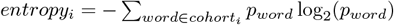

Word probabilities (*p*_*word*_) were taken from SUBTLEX-NL (Keuleers et al., 2010) and converted to phonemic transcriptions using the G2P service of Kisler et al. (2017) (using the specifications: lex output, x-sampa inventory, and language Dutch). This dictionary allowed us to determine the active cohort at each phoneme position.

Cohort entropy was computed for all phonemes except the initial phoneme of each word. At word onset, entropy reflects only the distribution of words beginning with that phoneme, not competition reduction over time. For analysis, non-onset phonemes were divided into low- and high-entropy categories: phonemes below the 33rd percentile of entropy values were labeled *low-entropy*, and those above the 66th percentile were labeled *high-entropy*. Note that, for this analysis, roughly half of the phonemes were discarded, as word-onset phonemes and those with entropy values between the 33rd and 66th percentiles were excluded. An overview of the occurrences of each phoneme descriptor can be found in Figure S2.

### 2.5 EEG preprocessing

EEG preprocessing was performed in Python (version 3.11.7) using MNE-Python (Gramfort et al., 2013). Preprocessing followed a two-step approach: (1) artifact identification on the concatenated recordings to ensure consistency across the multiple story parts, and (2) story-part-specific preprocessing using this artifact information from step 1.

#### Step 1: Artifact identification on concatenated data

To minimize the bias of artifacts on the components identified with independent component analysis (ICA), the four story segments presented in silence were concatenated into one continuous recording.

Artifactual channels were identified using the PREP pipeline (Bigdely-Shamlo et al., 2015) and excluded. The data were band-pass filtered between 1–40 Hz. To detect transient artifacts, a custom amplitude-based thresholding method was applied: time points where the smoothed signal exceeded ten standard deviations were flagged, together with a 2 second margin before and after. The threshold of ten standard deviations was chosen arbitrarily based on visual inspection of the data, ensuring that the largest artifacts, primarily those caused by muscle activity and motion, were flagged. These segments were removed prior to ICA to prevent large artifacts from biasing the component estimation. ICA was performed on the cleaned dataset using the infomax algorithm, decomposing the data into 30 independent components. The number of independent components was set arbitrarily, ensuring it was sufficiently high to capture artifacts within distinct components while minimizing the inclusion of neural activity within these artifact-related components. Artifact-related components (e.g., blinks, muscle activity, heartbeat; on average, 8 *±* 4 artifactual components) were manually identified and marked for removal.

#### Step 2: Segment-specific preprocessing

Each story segment was then processed individually, applying the artifact information from Step 1.

Previously identified bad channels were removed, and the recordings were cropped to the time window of speech presentation. Data were downsampled to 256 Hz. Artifact-related components identified in Step 1 were removed using ICA back-projection. Signals were re-referenced to the common average and band-pass filtered between 0.5–25 Hz. Bad channels were interpolated. A final downsampling to 128 Hz was applied, followed by parametric normalization per channel.

#### Epoching

The cleaned EEG data were segmented into epochs time-locked to phoneme onsets, ranging from 200 ms before to 600 ms after each onset.

### 2.6 Neural dynamics of phoneme processing: Temporal Generalization approach

We assessed the neural dynamics of phoneme processing using temporal generalization decoding (Gwilliams et al., 2022). Using this approach, we derived 3 measures:

#### 1. Temporal generalization decoding performance

refers to the 2D matrix that illustrates how accurately a phonetic descriptor can be decoded across all combinations of training and testing times. This measure indicates when a phonetic descriptor can be decoded from the neural response, thereby providing insight into when a phoneme is represented in the brain. Examining this matrix enables inferences about the stability of neural phonetic representations.

#### 2. Decoding duration

reflects the decoding performance along the diagonal of the 2D temporal generalization matrix, i.e., when training and testing times are identical. The period during which this decoding performance remains significantly above chance level provides insight into how long the phoneme representation is maintained in verbal short-term memory.

#### 3. Decoding generalization

captures the width of the diagonal “finger,” which is the most common pattern observed in the 2D temporal generalization matrix. This measure reflects how long the same neural representation remains active and, consequently, quantifies the rate at which phonetic information propagates across neural populations, thereby indicating the speed of phoneme processing.

#### Decoding a phoneme descriptor from the neural response

For each phoneme descriptor (see Section 2.4), we trained a logistic regression classifier to predict whether the descriptor was present (*true*) or absent (*f alse*) from the EEG data at each time point in the phoneme-locked epoch (-200 to 600 ms; 1*× n*_*channels*_). Hence, 104 classifiers were estimated to cover the range from -200 to 600 ms (in steps of 7.8 ms, according to the 128 Hz sampling rate) when the classifier is trained and tested on the same time point in an epoch.

To train and test the classifier, we used 5-fold cross-validation. In each iteration, four folds were used for training, allowing the classifier to learn how EEG channel activations relate to the presence of a given phonetic descriptor. This learning process is thought to capture the neural activation patterns associated with the occurrence of that descriptor, which reflect the underlying neural source activity, i.e., the neural representation of the phoneme. Classifier training employed ℓ_2_-regularized logistic regression with a regularization parameter of 0.1, selected on a subset of the data.

Performance was assessed on the held-out testing fold, producing probability estimates for each test epoch, i.e., the likelihood that the observed EEG channel activations reflected the presence of the phoneme descriptor. These probabilities were then compared against the true labels to compute the area under the receiver operating characteristic curve (AUC), which served as the measure of decoding accuracy.

Notice that as the logistic regression classifier performs a binary classification: either the phoneme descriptor was present (*true*) or absent (*f alse*) from the EEG data at a specific time point in the phoneme-locked epoch. Hence, the chance level of the classifier is 0.5. The classifier’s performance, although often significant, marginally improves the classification accuracy above chance. Increasing the amount of data and subjects would enhance the classifier’s performance.

#### Temporal generalization

In temporal generalization decoding, the training and testing time points do not need to coincide. Instead, the classifier is trained on neural activity at one time point after phoneme onset and evaluated on activity from a different time point. For example, a classifier trained at 200 ms after phoneme onset can be tested at 100 ms relative to phoneme onset. This approach probes whether the neural representation learned at one time point generalizes to another, thereby revealing the temporal stability of the underlying representation. Repeating this procedure across all train–test pairs (10816 possible combinations) produced a two-dimensional temporal generalization matrix spanning the -200 to 600 ms window.

#### Quantifying the temporal generalization matrix

Two complementary aspects of the matrix were analyzed:

- **Decoding duration**: Defined along the diagonal of the temporal generalization matrix, where training and testing occur at the same time point. This measure reflects the duration for which phonetic descriptors can be decoded from the EEG channel activations. Conceptually, it indicates the time window over which a phoneme is actively processed by the brain and may therefore capture the extent to which a phoneme is maintained in verbal short-term memory.
- **Decoding generalization**: Defined by the off-diagonal values, or equivalently by the width of the diagonal, where training and testing occur at different time points. This measure reflects the temporal stability of neural representations, i.e., how the neural representation at one time point generalizes to another time point. A static representation produces a block of significant decoding across a time window, whereas a dynamic representation yields significance primarily along the diagonal. Thus, decoding generalization captures how long a neural phoneme representation remains active within the same neural code configuration, allowing inferences about the speed at which phonemes are transferred from one neural population to another. To quantify decoding generalization, the two-dimensional matrix was rotated and shifted such that the diagonal corresponded to 0 ms (equal training–testing times). The data were then averaged across training times (y-axis) and z-scored, yielding a bell-shaped generalization curve. The width of this curve at 20% of its maximum amplitude was defined as the duration of neural generalization. Specifically, the analysis was restricted to a *±* 200 ms training–testing difference, with the left and right sides of the curve around the peak considered separately. Linear interpolation was then used to identify the points at which the curve crossed 20% of the peak value, and the difference between these two crossings was taken as the duration of generalization. The generalization duration is a marker indicating how long a specific neural representation of a phonetic descriptor remains temporally stable. Hence, it suggests a measure of processing speed: the longer the generalization duration, the less dynamic and slower the phoneme processing. Because of rotation, averaging across the y-axis and z-scoring the curve amplitude are not interpretable. Therefore, for visualization purposes, the curves are rescaled to share the same maximum value.

#### Spatial decoding performance

To examine the spatial distribution of decoding performance, classifiers were trained separately on each EEG channel, yielding a scalp-wide map of decoding performance. For computational efficiency, the full epoch time course was used as input when decoding the phonetic descriptor. Because the entire time course served as input to the classifier, temporal information was not preserved. Consequently, no inferences can be made about which time points contributed to the observed differences in spatial decoding performance.

### 2.7 Overview of performed analysis

A specific set of analyses was conducted to address each research aim.

- **Feasibility of decoding phoneme processing in children**
  - **Data**: All 25 children (irrespective of dyslexia diagnosis).
  - **Analyses**:
    * Decoding performance of individual phonetic descriptors (all phoneme epochs).
    * Temporal generalization: significance of decoding duration and decoding generalization, averaged across descriptors (all phoneme epochs).
    * Comparison of word-onset vs. non-onset phonemes.
    * Comparison of low-vs. high-entropy phonemes (non-onset phoneme epochs only).
- **Group differences: children with vs. without dyslexia**
  - **Data**: Age-matched subgroups of 9 children with dyslexia and 9 age-matched typical readers.
  - **Analyses**:
    * Group comparison of phoneme decoding (including decoding duration and generalization), repeated for: all phoneme epochs, word-onset phonemes only, non-onset phonemes only.
    * Group comparison of low-vs. high-entropy phonemes (non-onset phonemes).
    * Differences in spatial decoding performance (only reported when a significant difference was observed)
- **Individual differences: relation to reading-related behavioral measures**
  - **Data**: All 25 children (irrespective of dyslexia diagnosis).
  - **Analyses**:
    * Correlations between reading-related behavioral measures and decoding duration, averaged across descriptors.
    * Analyses repeated for: word-onset phonemes only, non-onset phonemes only, and (non-onset) low-entropy phonemes only.

### 2.8 Statistics

Within groups, decoding performance was tested against baseline using one-sample cluster-based permutation tests (MNE-Python; Gramfort et al., 2013). For both the 2D temporal generalization matrix and the derived decoding duration, the chance level corresponded to 0.5. To center this data at zero under the null hypothesis, 0.5 was subtracted prior to testing.

Between groups, two-sample cluster-based permutation tests were applied. All cluster tests (one-sample and two-sample) were run with 1000 permutations, using a cluster-forming threshold corresponding to *p* = 0.05. For multiple comparisons (e.g., separate tests for dyslexic vs. typical readers), the threshold was Bonferroni-corrected (e.g., *p* = 0.05*/*2). If test results appeared unstable, the number of permutations was increased to 3000 to improve robustness.

To assess group differences in the duration of decoding generalization (i.e., the width of the bell-shaped curve at 20% of maximum amplitude), we used a two-sided Mann–Whitney U test (SciPy toolkit; (Jones et al., 2001)).

## 3 Results

### 3.1 Neural Dynamics of Phoneme Processing in Children

#### 3.1.1 Phonetic Descriptors Are Encoded in Children’s EEG Data

Previous studies have shown that phoneme categories can be decoded from neural responses. Gwilliams et al. (2022) demonstrated successful decoding from MEG recordings in adults listening to two hours of speech, particularly within 50–300 ms after phoneme onset. Similarly, Kries et al. (2024) reported that phonetic features could be decoded from EEG responses during 25 minutes of story listening in individuals with aphasia. However, this approach has not yet been validated in young children.

Figure 2, panel A, illustrates the decoding time course, reflecting the extent to which phoneme categories can be extracted from EEG responses. A one-sample cluster-based permutation test (*p*_corrected_ = 0.05 across 18 features) revealed that most phonetic descriptors are decodable above chance (Panel A), except the *labial* phonetic descriptor concerning the place of articulation, and *back* phonetic descriptor regarding tongue front-backness. Consistent with prior findings (Gwilliams et al., 2022; Kries et al., 2024), significant decoding typically occurs between 50 and 300 ms post-onset (see Table S1 for timing details). The average peak of maximal decoding performance, if a cluster was observed between 0 and 300 ms, was situated around 154 ms.

**Figure 2.**
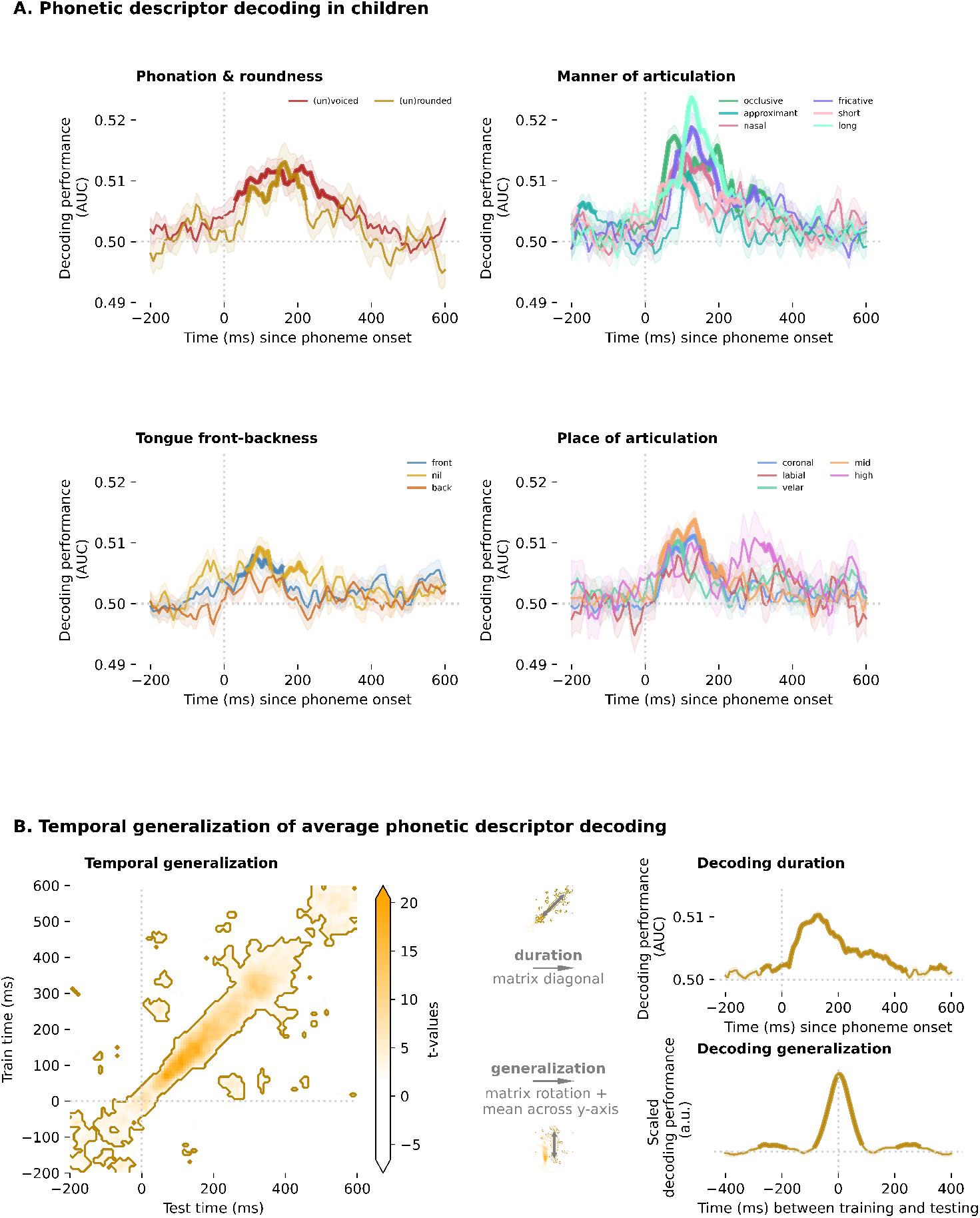
Neural dynamics of phoneme processing in children. **Panel A**: Above-chance decoding of 18 phonetic descriptors when training and testing times are the same, i.e., the decoding duration. The thicker segments indicate statistically significant time points. It should be noted that when pronunciation classes are limited to only two phoneme descriptors (i.e., phonation and roundness), the decoding performance of these descriptors is identical. This occurs because each phoneme is classified into one of two mutually exclusive categories, e.g., voiced versus unvoiced, leading to perfectly mirrored labels and, consequently, equivalent decoding performance. **Panel B**: Temporal generalization of the average phonetic descriptor across train-test time combinations (left). Significant decoding clusters are outlined. Decoding duration (top right) and generalization (bottom right) are also shown, with thicker segments marking significant intervals.

Due to variability in duration and occurrence across descriptors, decoding performance is often averaged across features, as shown in Panel B. The average decoding performance of the phonetic descriptor was significantly above chance, with the main diagonal cluster spanning from -130 to 467 ms (*p* = 0.001; average decoding = 0.503; Figure 2.B left panel). For decoding duration, three significant clusters were identified, the largest extending from -29 to 437 ms (*p <* 0.001; average decoding = 0.505; Figure 2.B top right panel). Additionally, decoding generalization was significantly greater than zero, suggesting that neural representations of phonetic features are temporally stable across an average window of 149 ms (Figure 2.B bottom right panel).

#### 3.1.2 Word-onset phonemes are encoded differently in a child’s brain compared to non-onset phonemes

Subsequently, we asked whether word-onset phonemes are processed differently by the brain compared to non-onset phonemes. Word-onset phonemes may capture not only phonemic encoding but also word-initial and anticipatory effects, since word-initial phonemes play a crucial role in word recognition because they strongly constrain the set of activated lexical candidates. Hence, they play an important role in the top-down process of generating predictions on the upcoming sensory input. Moreover, individuals with dyslexia show lower synchronization to simple sound fragments modulated at word frequency (De Vos et al., 2017, 4 Hz), suggesting also challenges

in processing bottom-up acoustic sensory input at the word rate. This raises the possibility that word-onset phonemes may be processed differently. Before comparing children with and without dyslexia, however, we first examined whether phoneme position, word-onset versus non-onset, affects neural encoding independently of dyslexia.

Our results indicate that word-onset and non-onset phonemes are processed differently. As shown by the black, dashed contours in Figure 3.A, significant differences in neural dynamics were observed from -200 to 227 ms (*p* = 0.002) and from -200 to 234 ms (*p* = 0.003), with decoding performance for word-onset phonemes exceeding that of non-onset phonemes (average difference: 0.003 and 0.004, per cluster, respectively). This suggests that the representation of word-onset phonemes can be decoded from the neural data prior to word onset.

**Figure 3.**
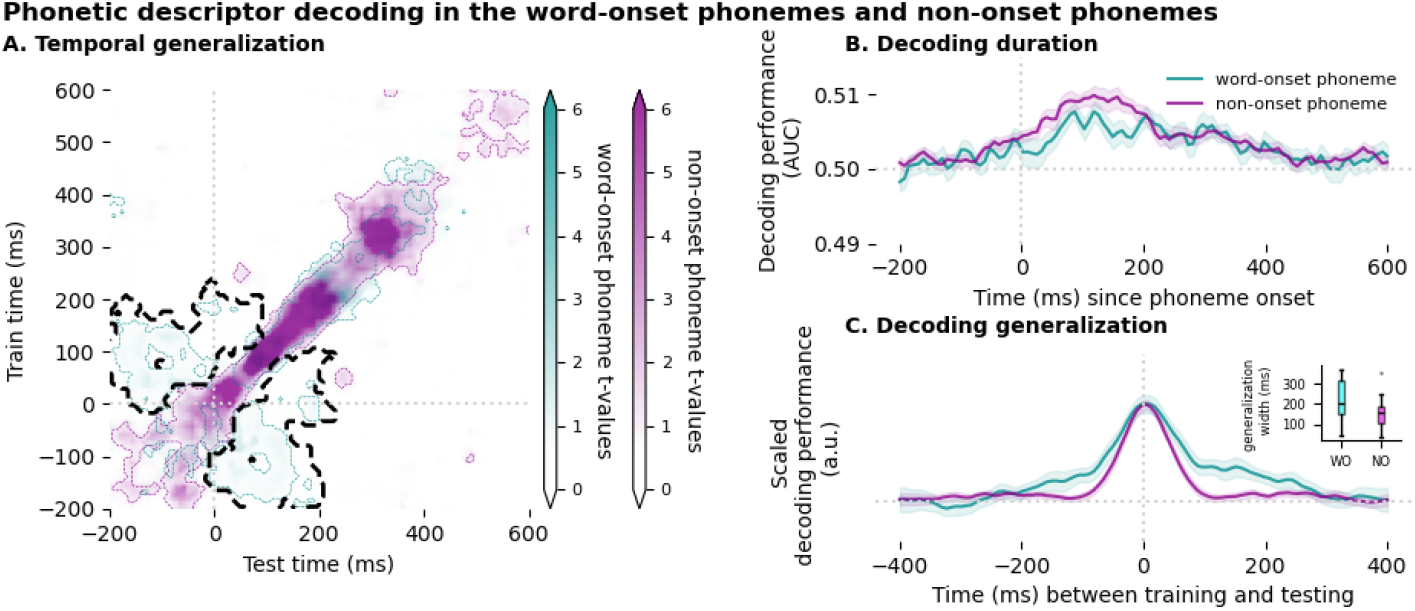
Neural dynamics of word-onset and non-onset phoneme processing. **Panel A**: Temporal generalization of the average phonetic descriptor for word-onset (teal) and non-onset (magenta) phonemes across train-test time combinations. Colored contours indicate clusters with decoding performance significantly above chance. The bold, striped black contour highlights clusters where neural dynamics significantly differ between conditions. **Panel B**: No significant difference in decoding duration between word-onset and non-onset phonemes. **Panel C**: Decoding generalization curves for word-onset (WO) and non-onset (NO) phonemes. The inset shows the difference in generalization duration.

Despite these differences, decoding duration did not significantly differ between word-onset and non-onset phonemes (Figure 3.B). The width of the bell-shaped decoding generalization curves (Figure 3.C), reflecting the temporal stability of phonetic representations, did not differ significantly. Generalization lasted on average 214 ms for word-onset phonemes and 155 ms for non-onset phonemes (Mann–Whitney U-test; *p* = 0.07).

#### 3.1.3 Neural dynamics of low and high entropy phonemes are different

One can expect that high- and low-entropy phonemes are processed differently. High-entropy phonemes are associated with greater lexical competition, meaning that many potential lexical candidates remain active. Consequently, predicting the upcoming phoneme becomes more challenging. In contrast, low-entropy phonemes, being more predictable, likely engage stronger top-down linguistic processes. Indeed, Gwilliams et al. (2022) and Kries et al. (2024) demonstrated that phonemes are processed differently depending on their lexical competition. Specifically, phonemes with high entropy, reflecting strong competition, as many words remain possible, exhibit higher decoding performance in later time windows after phoneme onset (300–330 ms) compared to phonemes with low entropy, which face weaker competition. Taken together, these findings suggest that high-entropy phonemes, characterized by greater lexical uncertainty, can be decoded for a longer period, indicating that they are maintained in neural processing for a longer duration.

Unexpectedly, no such difference was found in the overall neural dynamics of high-versus low-entropy phoneme processing when assessed using the 2D temporal generalization matrix (Figure 4). However, analysis of the diagonal decoding performance revealed a significant cluster from 452 to 530 ms (*p* = 0.036), indicating that low-entropy phonemes were decoded with higher accuracy than high-entropy phonemes. This trend contrasts with findings from Gwilliams et al. (2022) in adults and suggests that low-entropy phonemes may be maintained longer in neural representations in children, rather than high-entropy phonemes.

**Figure 4.**
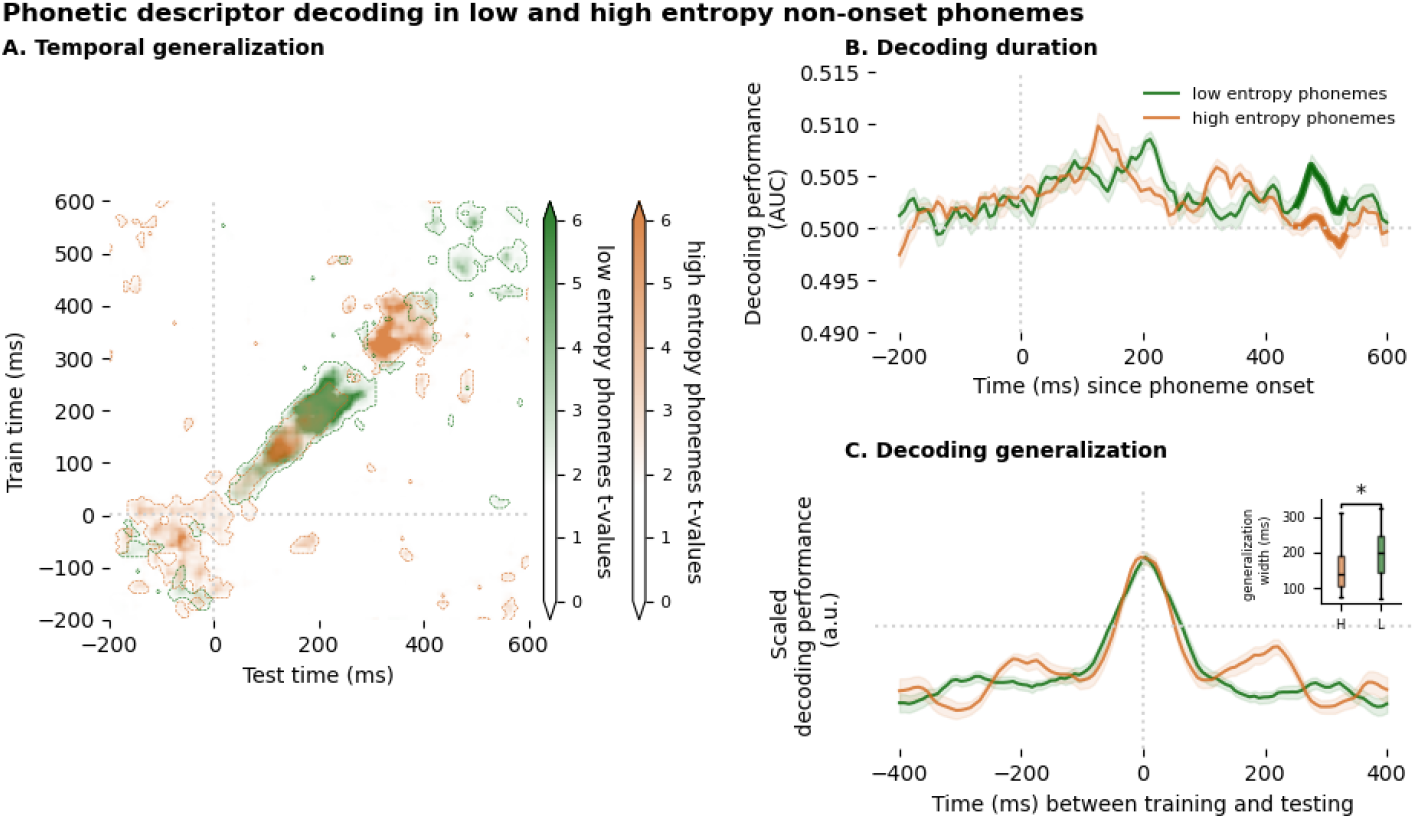
Neural dynamics of low- and high-entropy non-onset phoneme processing. **Panel A**: Temporal generalization of the average phonetic descriptor for low-entropy (green) and high-entropy (orange) phonemes across train-test time combinations. Colored contours indicate clusters with decoding performance significantly above chance. No significant difference in overall neural dynamics was observed between conditions. **Panel B**: Decoding duration did not significantly differ between low- and high-entropy phonemes. **Panel C**: Decoding generalization differed between conditions. Thicker line segments mark time points with significant differences in average decoding performance.

The generalization duration differed significantly: low-entropy phonemes generalized over a longer period (197 ms) compared to high-entropy phonemes (153 ms; Mann–Whitney U-test; *p* = 0.039).

### 3.2 Altered neural dynamics of phoneme processing in children with dyslexia

As investigating the neural dynamics in children is feasible, we examined whether differences are observed between children with dyslexia and their age-matched typical-reading peers.

#### 3.2.1 Altered temporal generalization in children with dyslexia

Comparing decoding performance across **all phonemes**, a cluster-based permutation test revealed a significant difference between typically reading children and children with dyslexia during later stages of phoneme processing (149–483 ms; Figure 5.1A; *p <* 0.01). Within this time window, children with dyslexia exhibited higher decoding accuracy than their typically reading peers. However, no significant group differences were found in decoding duration or decoding generalization (Figures 5.2A and 5.3A). The decoding duration differed significantly from chance for both groups (Figure 5.2A), as annotated by the horizontal lines below the plot. For typical readers, the significant cluster lasted 396 ms (from -21 to 274 ms). For dyslexic readers, the cluster lasted 365 ms (from 33 to 398 ms). It is important to note that different statistical tests were performed: one comparing decoding performance between the two groups, and others comparing decoding performance to chance level per group. Because the degrees of freedom and the corrections for multiple comparisons differ between these tests, their results are not necessarily expected to converge. Spatially, decoding performance differed in central left-hemispheric regions (*p* = 0.047), specifically at channels *CP5, CP3*, and *CP1*, where children with dyslexia showed higher phoneme encoding (Figure 5.4A).

**Figure 5.**
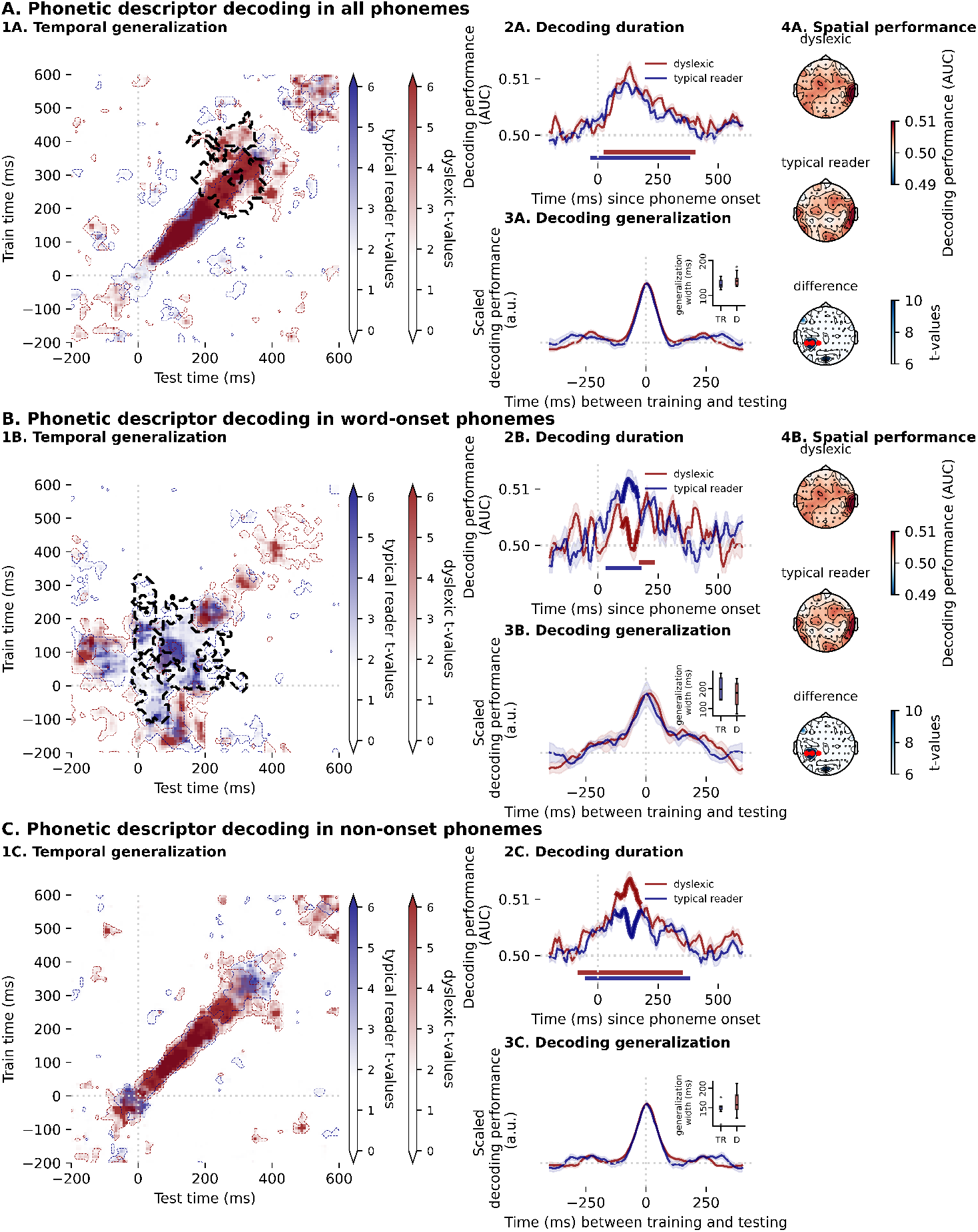
Neural dynamics of phoneme processing for all phonemes (**Panel A**), word-onset phonemes (**Panel B**), and non-onset phonemes (**Panel C**). **Subpanel 1**: Temporal generalization of the average phonetic descriptor in typical readers (blue) and children with dyslexia (red), shown across train-test time combinations. Colored contours indicate clusters with decoding performance significantly above chance; bold, striped black contours mark significant group differences. **Subpanel 2**: Decoding duration for both groups, with thicker lines indicating time points of significant differences. The horizontal line below the plot, in the respective color, indicates whenever the decoding performance is significantly above chance. **Subpanel 3**: No significant group differences in decoding generalization. **Subpanel 4**: Spatial decoding differences were observed in right-hemispheric regions for all phonemes (Panel A) and word-onset phonemes (Panel B), with significant channels annotated in red. No spatial differences were found for non-onset phonemes.

As shown in Section 3.1.2, word-onset phonemes are encoded differently from non-onset phonemes in children’s neural responses. To further investigate group differences, we replicated the analysis separately for word-onset and non-onset phonemes. For **word-onset phonemes**, a significant group difference was observed in the 2D temporal generalization matrix between typical readers and children with dyslexia, occurring during early neural responses (–106 to 328 ms; *p <* 0.01; Figure 5.1B). During this time period, typical readers showed higher decoding performance. The same trend is seen for the decoding duration, which also differed significantly (102–157 ms; *p* = 0.019). The decoding duration differed significantly from chance for both groups (Figure 5.2B), as annotated by the horizontal lines below the plot. For typical readers, the significant cluster lasted 132 ms (from 40 to 172 ms). For dyslexic readers, the cluster lasted 46 ms (from 180 to 227 ms). No differences were found in the decoding generalization (Figure 5.3B). Spatially, a significant cluster was observed in central left-hemispheric regions (*CP5, CP3, CP1*; *p* = 0.037; Figure 5.4B), with children with dyslexia having higher decoding performance. The latter shows an effect opposite to that observed for decoding duration. Note that spatial decoding was assessed using the full epoch; therefore, temporal information was not preserved. As a result, later time points (around 400 ms) may have driven the observed spatial differences.

For **non-onset phonemes**, no significant group differences were found in the 2D temporal generalization matrix (Figure 5.1C). However, decoding duration differed significantly between groups (79–172 ms; *p* = 0.002; dyslexics obtained higher performance; Figure 5.2C). The decoding duration differed significantly from chance for both groups (Figure 5.2C), as annotated by the horizontal lines below the plot. For typical readers, the significant cluster lasted, in total, 403 ms (from -44 to 234 ms and from 250 to 374 ms). For dyslexic readers, the cluster lasted 419 ms (from -75 to 343 ms). The decoding generalization (Figure 5.3C) and spatial performance showed no significant differences.

Overall, when comparing the time windows during which decoding performance is significantly above chance (subpanel 2 in panels A, B and C), we observe that whenever the word-onset phoneme is included, i.e., for all phonemes or for word-onset phonemes only, typical readers exhibit a longer significant cluster. This suggests that word-onset phonemes may be maintained longer in the verbal short-term memory of typical readers compared to children with dyslexia. In contrast, the opposite pattern is observed for non-onset phonemes, where children with dyslexia show a longer duration of above-chance decoding, suggesting that non-onset phonemes are maintained longer in their verbal short-term memory.

Comparing panel B and C of Figure 5, it is clear that the decoding duration pattern in children with dyslexia is different, suggesting children with dyslexia process word-onset phonemes differently compared to non-onset phonemes. Indeed, when doing this direct comparison (see Figure S3), a significant difference is observed, which is not observed for their typical reading peers.

#### 3.2.2 Low entropy phonemes are encoded differently in a dyslexic child’s brain

As previously shown (Figure 4), high- and low-entropy phonemes are processed differently in children’s brains. We therefore examined whether these differences vary between typical readers and children with dyslexia.

Contrary to our expectations, no significant group differences were found in the processing of high-entropy phonemes, neither in the 2D temporal generalization matrix, decoding duration, decoding generalization, nor spatial performance. However, the decoding duration differed significantly from chance for both groups (Figure 6.2A), as annotated by the horizontal lines below the plot. For typical readers, the significant cluster lasted 69 ms (from 304 to 374 ms). For dyslexic readers, the cluster lasted 209 ms (from 25 to 234 ms).

**Figure 6.**
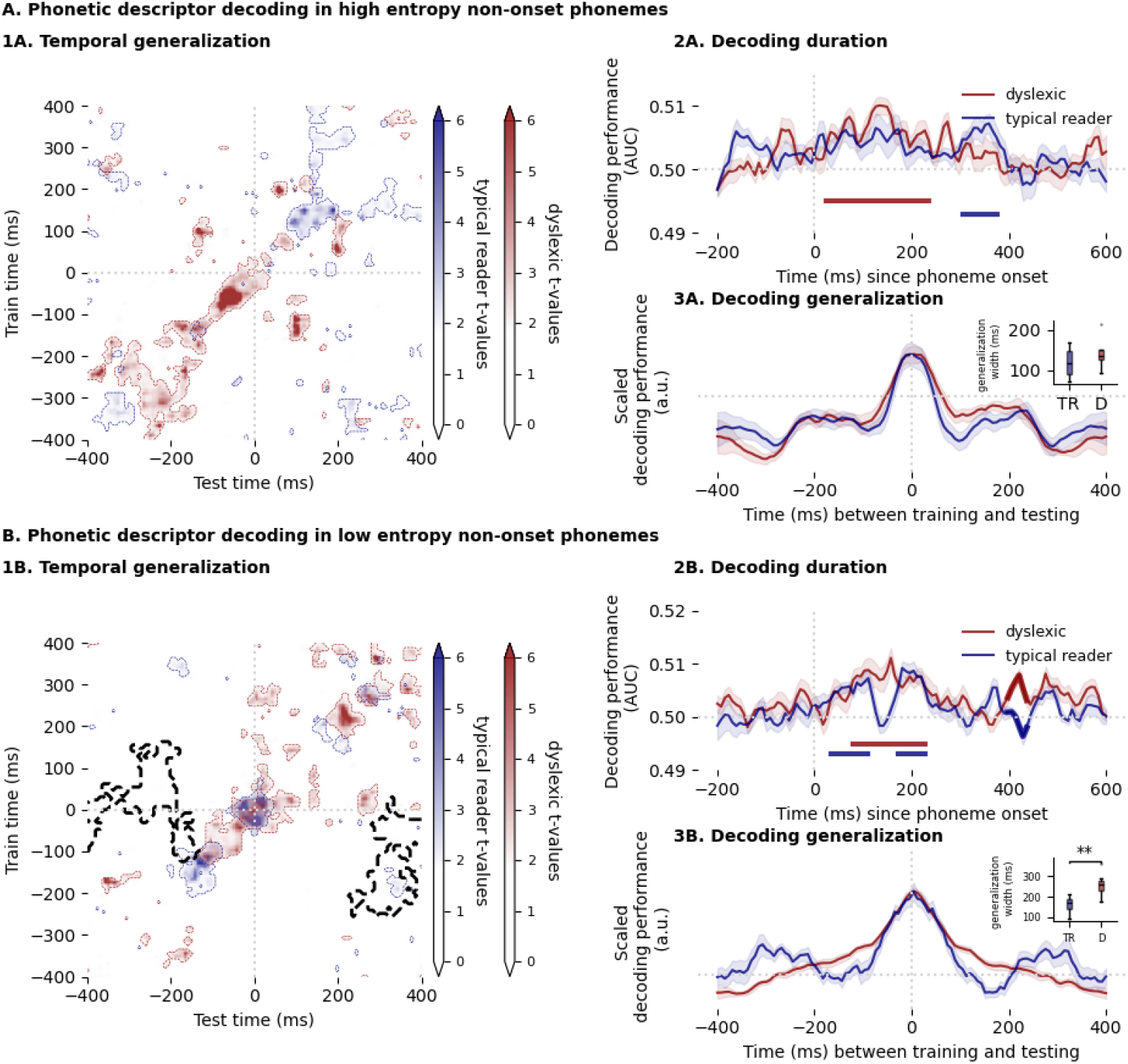
Neural dynamics of low- and high-entropy non-onset phoneme processing in typical readers and children with dyslexia. **Panel 1A**: Temporal generalization of high-entropy phonemes in typical readers (blue) and children with dyslexia (red), shown across train-test time combinations. Colored contours indicate clusters with decoding performance significantly above chance. No significant group differences were observed. **Panel 2A**: Decoding duration for high-entropy phonemes did not differ significantly between groups. The horizontal line below the plot, in the respective color, indicates whenever the decoding performance is significantly above chance. **Panel 3A**: No significant differences in decoding generalization for high-entropy phonemes. **Panel 1B**: Temporal generalization of low-entropy phonemes in typical readers and children with dyslexia. Colored contours indicate above-chance decoding; the black, striped contour marks significant group differences. **Panel 2B**: Decoding duration differed significantly between groups, with thicker line segments indicating significant time points. The horizontal line below the plot, in the respective color, indicates whenever the decoding performance is significantly above chance. **Panel 3B**: Decoding generalization also differed significantly. The inset highlights that children with dyslexia showed longer neural generalization than typical readers.

For low-entropy phonemes, however, two significant clusters emerged in the temporal generalization matrix: one from –52 to 600 ms (*p* = 0.020), and another from –200 to 359 ms (*p* = 0.032), with typical readers showing slightly higher decoding performance (see Figure 6.1B). In decoding duration, a late cluster from 398 to 436 ms (*p* = 0.035) revealed that the classifier obtained a better performance with the neural data of children with dyslexia (average difference: 0.006). The decoding duration differed significantly from chance for both groups (Figure 6.2B), as annotated by the horizontal lines below the plot. For typical readers, the significant cluster lasted, in total, 131 ms (from 33 to 110 ms and from 172 to 227 ms). For dyslexic readers, the cluster lasted 147 ms (from 79 to 227 ms). When quantifying generalization duration, children with dyslexia exhibited longer neural generalization (258 ms) compared to typical readers (164 ms), with a group difference of 94 ms (Mann–Whitney U-test; *p* = 0.006). No significant group differences were found in the processing of low-entropy phonemes for the decoding generalization, nor in spatial performance.

When comparing the duration of clusters where decoding performance is significantly above chance (subpanel 2 in panels A and B), we observe that for both low- and high-entropy non-onset phonemes, children with dyslexia exhibit longer durations. This suggests that non-onset phonemes are maintained longer in their verbal short-term memory, independent of their lexical relevance. Interestingly, for high-entropy phonemes, the location of this cluster differs markedly: typical readers show a late cluster around 350 ms, whereas around this time, no significant decoding performance is observed for children with dyslexia.

### 3.3 Link between neural dynamics of phoneme processing and reading-related behavioral measures

To quantify the relationship between neural dynamics of phoneme processing and reading-related behavioral measures, we correlated reading-related behavioral measures with the diagonal of the 2D temporal generalization matrix, referred to as decoding duration, at each time point. This analysis was conducted for word-onset phonemes, non-onset phonemes, and low-entropy phonemes, as these categories previously showed the most prominent group differences between typical readers and children with dyslexia. All children were included in this analysis, encompassing both children with dyslexia and their age-matched typically reading peers.

Importantly, this was an exploratory analysis, and the observed effects were marginal, failing to survive correction for multiple comparisons. Nevertheless, as shown in Figure 7 and Table 1, the timing of these uncorrected significant correlations, around 150 and 450 ms, closely aligns with previously identified group differences in phoneme processing (see Figure 5, Panels B and C, for clusters near 150 ms; and Figure 6 for the cluster near 450 ms).

**Table 1:** Overview of the *uncorrected* significant correlations between the reading-related behavioral measures and the decoding duration.

| Reading-related behavioral measure | Phoneme type | Time window | Average correlation (spearman) |
| --- | --- | --- | --- |
| Rise time discrimination task | non-onset phonemes | 444 - 460 ms | -0.44 |
| CELF - phoneme section | word-onset phonemes | 133 - 165 ms | 0.49 |
| CELF - phoneme section | non-onset phonemes | 133 - 141 ms | -0.51 |
| CELF - phoneme section | non-onset phonemes | 452 - 460 ms | -0.45 |
| Phoneme deletion | word-onset phonemes | 405 - 421 ms | -0.57 |
| Phoneme deletion | word-onset phonemes | 460 - 475 ms | -0.46 |
| Phoneme deletion | non-onset phonemes | 126 - 149 ms | -0.48 |
| Phoneme deletion | low entropy phonemes | 95 - 110 ms | -0.52 |
| Phoneme deletion | low entropy phonemes | 250 - 258 ms | -0.45 |
| Phoneme deletion | low entropy phonemes | 398 - 405 ms | -0.49 |
| Phoneme deletion | low entropy phonemes | 421 - 429 ms | -0.49 |
| Non-word repetition task | word-onset phonemes | 56 - 64 ms | 0.44 |
| Non-word repetition task | low entropy phonemes | 250 - 258 ms | -0.48 |
| Non-word repetition task | low entropy phonemes | 398 - 429 ms | -0.51 |

**Figure 7.**
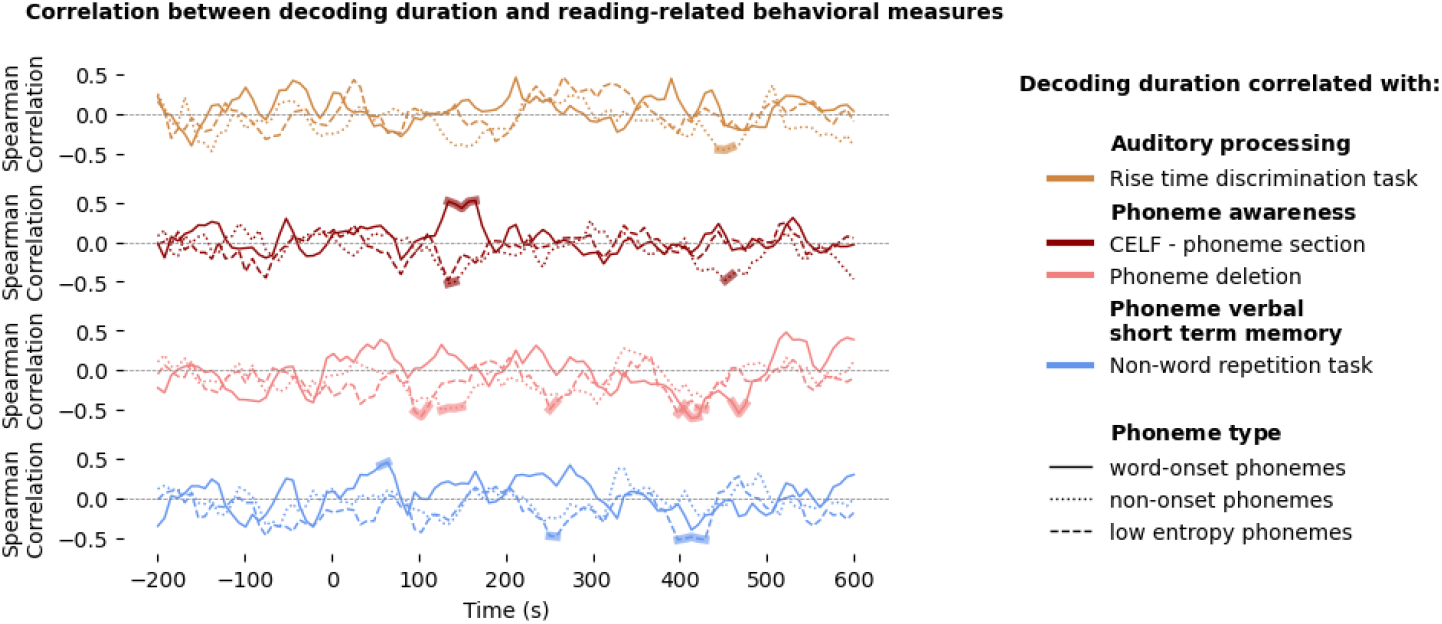
Correlation with the TG duration and the reading-related behavioral measures. The thinker line indicates whenever the correlation at this time point is significant; however, it is not corrected for multiple comparisons

Interestingly, the Spearman correlations at these time points were predominantly negative, suggesting that lower scores on reading-related behavioral measures are associated with higher decoding performance. An important remark is that observed correlations depend on the time window (the trends around 150 ms might differ from those observed in later processing around 400 ms) and phoneme analysis (word-onset, non-onset or low-entropy phoneme). This pattern may reflect compensatory neural mechanisms or altered processing strategies in children with weaker reading-related skills.

## 4 Discussion

This study evaluated the neural dynamics of underlying phoneme processing in children with and without dyslexia, when processing naturalistic continuous speech. In addition to quantifying the key differences in phoneme processing between typical readers and children with dyslexia, we explored the relationship between neural metrics of phoneme encoding and behavioral measures of reading-related behavioral measures, offering insight into the developmental connections between auditory processing and early literacy skills.

### 4.1 Neural dynamics of phonetic processing can be investigated in children

Our findings demonstrate that investigating the neural dynamics of phonetic processing in children is feasible, even with inherently noisier EEG data and limited recording durations. We successfully decoded individual phonetic descriptors from neural responses (Figure 2.A), with significant decoding observed between 70 and 200 ms post-phoneme onset, an average duration of 130 ms. This is notably shorter than the 300 ms window reported in adults (Gwilliams et al., 2022), likely reflecting developmental differences. At an age of seven, children’s phonemic processing remains in a dynamic phase of maturation, as evidenced by performance in behavioral contrast tasks (Hazan and Barrett, 2000) and neural envelope tracking during continuous speech perception (Bertels et al., 2023). Thus, the reduced decoding duration compared to adults may reflect the evolving nature of phoneme representation in the developing brain.

Interestingly, a higher decoding performance was observed on the early response to word-onset phonemes compared to non-onset phonemes (Figure 3). These findings may suggest the presence of anticipatory processing. Such processing is only feasible when the listener can reliably predict the upcoming word, implying that the brain engages in forecasting the next lexical item. Consequently, the observed effect may reflect a trace of predictive coding. The fact that word-onset phonemes exhibit anticipatory activity, whereas non-onset phonemes do not, could indicate that predictive mechanisms operate more strongly at the word level than at the phoneme level.

Alternatively, a purely acoustic explanation may be at play: word boundaries are often more clearly perceived than phoneme boundaries, as a brief silence typically precedes them. This pause may facilitate the observation of anticipatory processing at word onset, while anticipatory responses to subsequent phonemes may be obscured by the neural response to the preceding phoneme, making them more difficult to detect. Finally, non-onset phonemes tend to be less salient for word recognition and are more susceptible to coarticulatory influences, which can reduce their acoustic clarity and, in turn, the likelihood of predictive engagement.

In adults, Gwilliams et al. (2022) reported higher decoding performance for high-entropy phonemes around 300 ms and beyond 400 ms. In children, we did not observe a significant effect at 300 ms, though trends suggest higher decoding for high-entropy phonemes in that window. However, from 400 ms onward, we observed the opposite: low-entropy phonemes were decoded more accurately. While the cluster-based permutation test does allow identifying a shift in latency, the pattern may reflect the delayed processing of low-entropy phonemes (see Figure 6.2B). Such a delay is unexpected, as low-entropy phonemes are more predictable and, therefore, typically require less effortful processing. Nevertheless, low-entropy phonemes also showed longer duration of neural generalization (Figure 4), suggesting that the processing speed of these phonemes is lower compared to high-entropy phonemes despite contributing less linguistic novelty.

It is important to note that all children in this study were identified as at cognitive risk for dyslexia in kindergarten, which may bias results toward weaker reading profiles. Nonetheless, the temporal generalization approach showed feasibility in pediatric EEG data, paving the way for future research into phonological development across childhood.

### 4.2 Reduced early phonetic decoding performance to word-onset phonemes in children with dyslexia

A reduced decoding performance of the early neural response to word-onset phonemes is observed in children with dyslexia compared to typical readers (Figure 5.B). Word-onset phonemes play a crucial role in word recognition, as they strongly influence the activation of lexical candidates and are less affected by coarticulation. A clear perception of these initial phonemes is essential for successful word identification. The diminished early response in children with dyslexia may therefore hinder word recognition. Supporting this, Bonte and Blomert (2004) reported reduced ERP responses (120–240 ms) to target words in children with dyslexia and found anomalous contributions of phonetic/phonological cues, particularly word-onset information, to spoken word recognition. This may explain why individuals with dyslexia often struggle with speech comprehension in noisy environments (see review Calcus et al., 2018), where word-onset phonemes are more difficult to detect, potentially disrupting recognition of subsequent phonemes and the entire word.

We observed that word-onset phonemes can be decoded for a longer duration in typically reading children (Figure 5.2B) compared to children with dyslexia, suggesting that these phonemes are maintained longer in verbal short-term memory. If individuals with dyslexia relied more strongly on higher-level predictive coding mechanisms as a compensatory strategy, the opposite pattern would have been expected. Future research should therefore focus on a more thorough analysis of higher-level linguistic processing of words to clarify this effect. However, it is possible that the observed bottleneck is instead related to bottom-up acoustic processing.

In line with this interpretation, Van Hirtum et al. (2021) successfully evaluated an envelope enhancement strategy aimed at improving speech understanding in noise among children with dyslexia. This approach amplifies aspects of the bottom-up acoustic signal, specifically sharp amplitude rises, thereby enhancing speech intelligibility. By applying envelope enhancement, the children with dyslexia performed instantaneously as well as their typical reading peers. Interestingly, these sharp amplitude rise enhancements, which are beneficial to improve speech understanding in persons with dyslexia, occurred most commonly at the onset of the word.

Why envelope enhancement improves speech perception in noise in children with dyslexia might be explained by the temporal sampling framework proposed by Goswami (2011). This framework suggests that persons with dyslexia have difficulties with processing the rise and falls of the amplitude of the speech envelope, i.e., how the slow-temporal variations of the speech are modulated. By investigating the neural response to sinusoidal amplitude-modulated tones, one can assess these neural responses to these slow modulations, i.e., auditory steady-state responses (ASSR), rather than the complex speech envelope itself, which includes speech processing ranging from acoustic to linguistic processing. Indeed, investigating these ASSRs to low-frequency modulations as low as 2 and 4 Hz, individuals with dyslexia do show lower responses compared to their typical reading peers (e.g., Hämäläinen et al., 2012; Poelmans et al., 2012; De Vos et al., 2017). This nicely aligns with the reduced response observed to word-onset phonemes in the current study, as 4 Hz “word-rate” is a dominant frequency in the modulation of continuous speech (Varnet et al., 2017).

Studies on continuous speech further support this hypothesis. Di Liberto et al. (2018) found reduced neural tracking of the speech envelope in children with dyslexia across specific frequency bands. Destoky et al. (2022) reported similar findings, with effects most pronounced in babble noise conditions. Mandke et al. (2023) observed decreased coherence between the speech envelope and neural signals, while Araújo et al. (2024) reported reduced delta band power (2 Hz) in fronto-central regions. These studies consistently highlight differences in low-frequency neural processing (up to 8 Hz), suggesting that impaired encoding of word-onset phonemes in dyslexia may stem from deficits in tracking slow temporal modulations rather than phoneme-level encoding alone.

Regarding spatial decoding, we observed increased phonetic descriptor decoding in central left-hemispheric regions in children with dyslexia. However, findings across studies investigating neural speech tracking remain inconsistent. Di Liberto et al. (2018) reported reduced phoneme tracking primarily in right-hemispheric areas, while Destoky et al. (2022) found no significant spatial differences in envelope tracking, and Mandke et al. (2023) noted decreased coherence bilaterally. These discrepancies may reflect differences in language, methodology, and analytic approaches across studies.

### 4.3 Increased phonetic decoding performance when processing non-onset phonemes and low-entropy phonemes in children with dyslexia

Children with dyslexia exhibited higher decoding performance for non-onset phonemes between 79 and 172 ms compared to typical readers (Figure 5.2C). A similar pattern was observed when comparing word-onset and non-onset phonemes within the dyslexic group: decoding performance was significantly higher for non-onset phonemes between 126 and 165 ms (Figure S3.2B). This contrast was not present in typical readers, where the non-significant trend even suggested the opposite, word-onset phonemes showed slightly higher decoding performance (Figure S3.2A).

Additionally, low-entropy non-onset phonemes were decoded with greater accuracy between 398 and 436 ms post-onset and showed longer significant decoding above chance suggesting that these phonemes are maintained longer in the verbal short-term memory (Figure 6.2B). Moreover, the neural representations of low-entropy non-onset phonemes remained generalized for a longer duration suggesting slower phoneme processing in children with dyslexia (Figure 6.3B). Together, these findings suggest that children with dyslexia may rely more heavily on non-onset and low-entropy phonemes during speech processing, potentially reflecting compensatory mechanisms or altered temporal encoding strategies.

Since the division between word-onset and non-onset phonemes, or the division between high and low entropy phonemes, is not commonly done in dyslexia research, it is difficult to compare the current results to existing literature.

### 4.4 Phonemic representations in children with dyslexia: interpretations from current findings

A key methodological note in this study is that phonemes were linked to their phonetic descriptors. While phonemes represent the abstract building blocks of spoken language, each phoneme can correspond to multiple phones, the actual acoustic realizations. For instance, the English phoneme /t/ may be realized as [*t*] or [*t*^*h*^]. By associating phonemes with their pronunciation features, we acknowledge a simplification of the rich articulatory and acoustic variability inherent in speech. Nevertheless, as shown by King and Taylor (2000), classifying speech sounds into five key phonetic classes captures substantial variability, justifying the use of phonetic descriptor decoding as a proxy for phoneme-level processing.

Another important note is that by grouping phonemes into different phonetic descriptors, we are not investigating the neural response to a single phoneme, but to a more general property shared across different, yet somewhat similar, phonemes. Consequently, even though there are multiple acoustic variations within a single phoneme, and especially within phonemes grouped under the same phonetic descriptor class, these variations are all mapped to a similar phonetic representation. Therefore, this approach reflects neural mapping not at the level of individual phonemes, but in a more general, class-based manner.

How do these findings inform the debate on phonological representations in dyslexia? While our results do not directly reveal how phonemes are represented in the brain, they do shed light on differences in neural phoneme processing, differences that likely depend on the integrity and accessibility of phonological representations. If phonological representations were degraded or access to them impaired, we would expect delayed neural responses or reduced decoding performance in children with dyslexia. However, when analyzing phoneme processing independent of position (i.e., all phonemes together; Figure 5.A), no such clear group differences were observed in decoding duration or generalization. In fact, when differences did emerge in the temporal generalization matrix in later time windows, or in spatial decoding, children with dyslexia often showed higher performance.

This pattern becomes more pronounced when examining low-entropy non-onset phonemes (Figure 6.B). Here, children with dyslexia exhibited higher decoding performance in later stages post-phoneme onset, and their neural representations were longer generalized to other timepoints, indicating lower processing speed from one neural population to the other. These findings suggest that low-entropy non-onset phonemes, which are typically considered less informative for linguistic context, are processed more intensively by children with dyslexia. One interpretation is that children with dyslexia may apply less efficient internal weighting, failing to distinguish between linguistically relevant and irrelevant phonemes, leading to uniformly intensive processing. In challenging listening conditions, this lack of prioritization could become a bottleneck, impairing speech comprehension.

Alternatively, the increased processing of non-onset phonemes may reflect compensatory mechanisms. If children with dyslexia struggle to process earlier phonemes, particularly word-onset phonemes, as observed, then subsequent phonemes may require more effortful and prolonged neural engagement.

### 4.5 Strength of phonetic descriptor decoding correlates with reading-related behavioral measures

When examining differences in the neural dynamics of phoneme processing between children with and without dyslexia, two distinct temporal patterns emerged: a difference in neural dynamics around 150 ms and around 400 ms. These effects were most pronounced in word-onset phoneme processing, non-onset phoneme processing, and low-entropy non-onset phoneme processing. Accordingly, these categories were correlated with behavioral measures reflecting reading-related behavioral measures. Importantly, this analysis included all children in the sample, not only those with dyslexia and their age-matched peers, and was exploratory in nature. None of the observed correlations survived correction for multiple comparisons.

Nonetheless, even with a broader participant pool, decoding performance around the 150 ms and 400 ms, showed correlations with reading-related behavioral measures, suggesting that these neural time windows may be informative for understanding reading development. Most correlations were negative, indicating that higher decoding performance was associated with lower scores on reading-related behavioral measures. Notably, the longest stretch of significant correlations was observed between phoneme verbal short-term memory and low-entropy non-onset phoneme processing, indicating that the processing of predictable phonemes might play a role in the expression of reading difficulties.

While these findings should be interpreted cautiously due to their exploratory nature and limited statistical robustness, they suggest that investigating continuous speech processing, and particularly the neural encoding of phoneme features, in a larger and more diverse sample of children could yield valuable insights into the neural basis of reading development.

### 4.6 Limitations

This study measured EEG responses to continuous speech in children identified in kindergarten as having a cognitive risk for developing dyslexia. As a result, the sample does not reflect a representative population of children. Importantly, early cognitive risk does not guarantee continued risk or diagnosis by second grade. However, the group of typical readers may be biased toward children with slightly weaker reading skills compared to peers without early risk (Snowling and Melby-Lervåg, 2016). Nevertheless, the presence of significant group differences suggests that when using a control group without such early risk, stronger effects might be observed.

Another limitation concerns the classification of dyslexia. The criteria used in this study differ from the official diagnostic standards applied in Belgium. Additionally, no reading-level-matched control group was included. Consequently, it is difficult to disentangle whether the observed neural differences reflect the core neurobiological basis of dyslexia or are instead driven by differences in reading experience, which itself shapes brain development. Furthermore, the sample size of 25 children is insufficient to capture the full heterogeneity of dyslexia, thereby limiting the generalizability of the findings. Nevertheless, it is promising to see that differences in neural phoneme processing can be observed, even with a limited number of children.

External factors such as the COVID-19 pandemic also likely influenced reading development. School closures during first and second grade introduced variability in reading exposure, depending on each child’s home environment, which may have affected both behavioral and neural outcomes. Nevertheless, in a similar longitudinal cohort, Blockmans et al. (2025) reported that the phonological delay, presumably caused by the school closures, had disappeared by the start of third grade, when reading scores were collected to determine whether a child was classified as dyslexic. Therefore, we can expect that this classification remains robust, despite the potential consequences of COVID-19.

A methodological caveat is that the interpretations of phoneme-related neural dynamics may still reflect aspects of acoustic processing. Acoustic features were not explicitly controlled for, and thus some effects may be driven by low-level auditory processing differences. However, in the comparison of high- and low-entropy non-onset phonemes, the phoneme distributions were similar across high- and low-entropy phonemes, suggesting that these specific findings are less likely to be confounded by acoustic variability.

Research on dyslexia inherently faces challenges due to the disorder’s complexity and variability. Diagnostic criteria differ across countries, and language-specific assessments further complicate cross-study comparisons. As in this study, research inclusion criteria often diverge from clinical standards, contributing to variability in findings. To address these issues, future research should prioritize larger sample sizes and collaborative efforts across labs and countries. Harmonizing protocols and pooling data could help build robust, representative datasets that capture the multifaceted nature of dyslexia.

## Conclusion

This study demonstrates the feasibility of using temporal generalization analysis to investigate the neural dynamics of phoneme processing in children, even with the inherent challenges of noisy EEG data and limited sample sizes. By decoding phonetic descriptors from neural responses to continuous speech, we revealed meaningful differences in phoneme processing between typical readers and children with dyslexia, particularly in early and late time windows around 150 ms and 400 ms.

Our findings suggest that children with dyslexia exhibit altered neural encoding of word-onset and low-entropy non-onset phonemes, with evidence of prolonged and more intensive processing of less linguistically relevant phonemes. These differences may reflect compensatory mechanisms or inefficiencies in phonological weighting, which could contribute to difficulties in speech comprehension, particularly in noisy environments.

## Acknowledgements

We would like to thank Shauni Van Herck and Tilde Van Hirtum for their effort to collect the data. This data was part of a big, longitudinal design carried out by Shauni Van Herck, Femke Vanden Bempt, Maria Economou, Maaike Vandermosten, Jan Wouters and Pol Ghesquie`re. Hence, we express gratitude to them and all the children participating in the study.

## Supplementary Material

### S.1 Framing of the current subsample of participants in the longitudinal study of Van Herck et al. (2023)

This study uses data collected by Van Herck et al. (2023), as part of a longitudinal project, whereby children were followed from the pre-reading stage (∼5 years old; last year of kindergarten) to acquired reading skills (∼8 years old; 3rd grade), with behavioral and neural data collected throughout. None had a family history of dyslexia. A subset of participants also received an intervention. The original study focused mainly on temporal auditory processing using EEG (i.e., auditory steady-state responses). However, at one time point (2nd grade; ∼7 years old), a subset of children also listened to an audiobook. The current study analyzes these audiobook EEG data, which have not been examined in previous work. Thus, we rely on a subset of both the participants and the time points from Van Herck et al. (2023), specifically children tested at the beginning of 2nd grade who all belonged to the passive control group.

**Figure S1:**
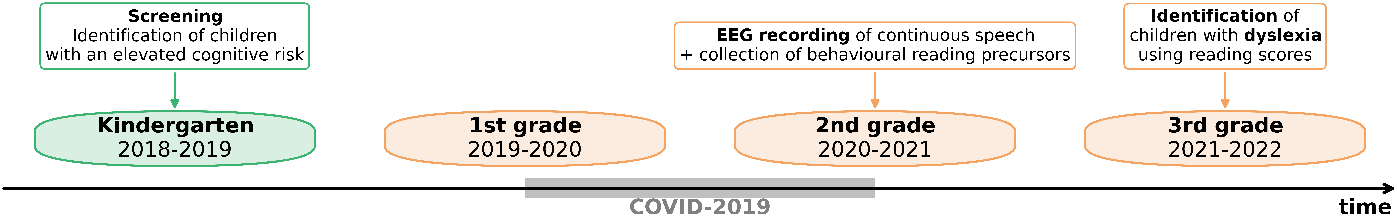
Overview of the data collection timeline.

### S.2 Phoneme Occurence

**Figure S2:**
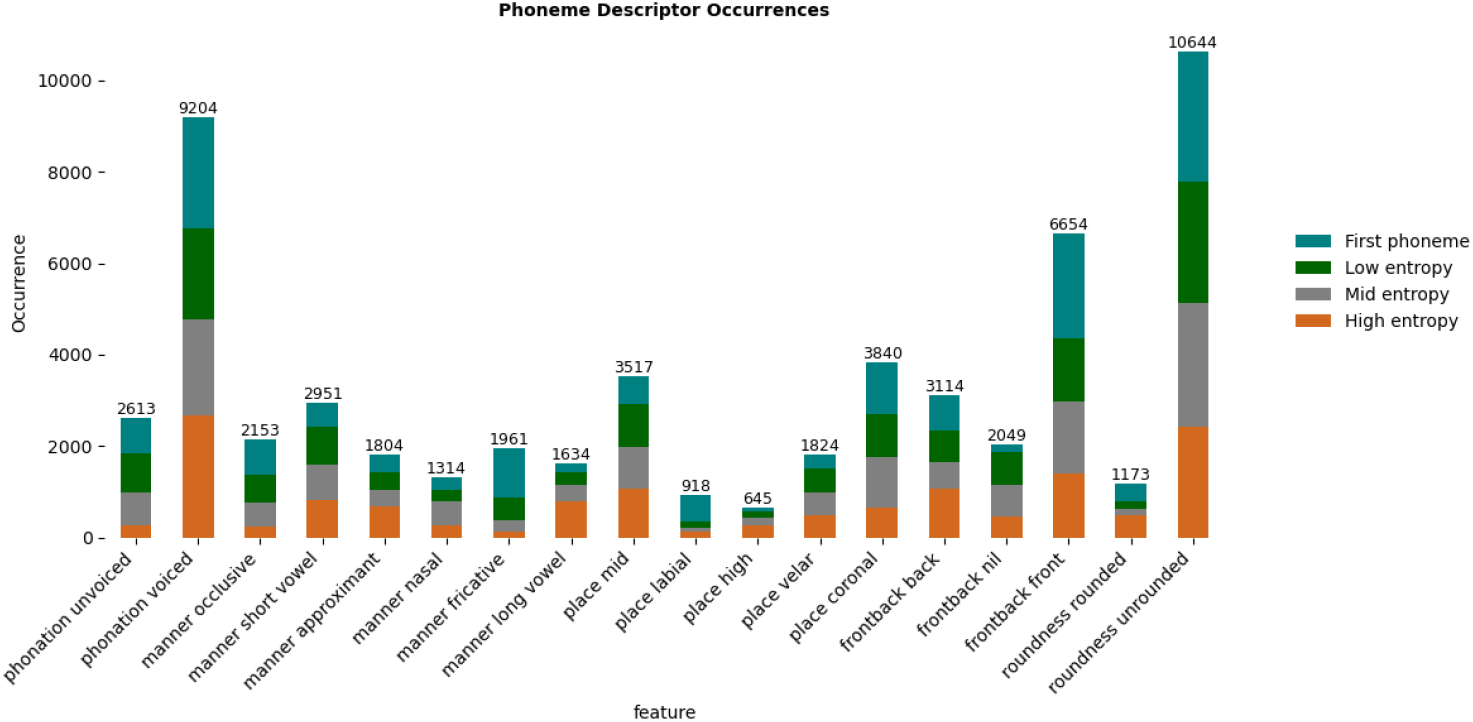
Overview of occurrence per phoneme descriptor.

### S.3 Above-chance decoding of phonetic descriptors in children

**Table S1:** Overview of the above-chance decoding of phonetic descriptors in children with its significant time window and the average decoding performance within this time window (*\*\*\* = p < 0*.*001, ** =p <0*.*01, * = p <0*.*05*)

| Pronunciation aspect | Descriptor | Significant decoding window | Average decoding performance | Significance |
| --- | --- | --- | --- | --- |
| phonation | (un)voiced | 33 - 320 ms | 0.510 | ** |
| roundness | (un)rounded | 64 - 219 ms | 0.509 | ** |
| manner | occlusive | 40 - 250 ms | 0.512 | ** |
| manner | occlusive | 281 - 343 ms | 0.508 | ** |
| manner | approximant | -176 - -137 ms | 0.506 | ** |
| manner | approximant | 102 - 141 ms | 0.509 | ** |
| manner | nasal | 56 - 180 ms | 0.512 | ** |
| manner | nasal | 234 - 266 ms | 0.508 | ** |
| manner | fricative | 71 - 211 ms | 0.513 | ** |
| manner | fricative | 234 - 304 ms | 0.507 | ** |
| manner | short vowel | 33 - 157 ms | 0.509 | ** |
| manner | short vowel | 188 - 266 ms | 0.506 | ** |
| manner | long vowel | 40 - 234 ms | 0.514 | ** |
| place | coronal | 48 - 196 ms | 0.508 | ** |
| place | velar | 48 - 110 ms | 0.508 | ** |
| place | mid | 40 - 211 ms | 0.509 | ** |
| place | high | 320 - 351 ms | 0.509 | ** |
| front-backness | front | 40 - 157 ms | 0.506 | ** |
| front-backness | nil | 64 - 126 ms | 0.507 | ** |
| front-backness | nil | 165 - 219 ms | 0.506 | ** |

### S.4 Children with dyslexia process word-onset phonemes differently from non-onset phonemes

Given that word-onset and non-onset phonemes are processed differently in children with dyslexia compared to typical readers, we investigated how these differences manifest within each group.

In both groups, neural dynamics diverged between word-onset and non-onset phoneme processing during early responses. Notably, most differences occurred outside the diagonal of the temporal generalization matrix, indicating that they do not align with identical train-test time points. In typical readers, a significant cluster was observed from –200 to 242 ms (*p* = 0.001), with word-onset phonemes decoded more accurately than non-onset phonemes (average difference: 0.006). No significant differences were found in decoding duration, generalization, generalization duration, or spatial distribution.

In children with dyslexia, two significant clusters emerged from –200 to 281 ms (*p* = 0.023) and –200 to 242 ms (*p* = 0.041), again showing higher decoding performance for word-onset phonemes (average difference: 0.005). However, decoding duration differed between conditions (126–165 ms; *p* = 0.024), with non-onset phonemes showing slightly higher performance (average difference: 0.011). A significant amplitude difference in decoding generalization was also observed (*p* = 0.006), though generalization duration and spatial decoding performance did not differ between phoneme types.

**Figure S3:**
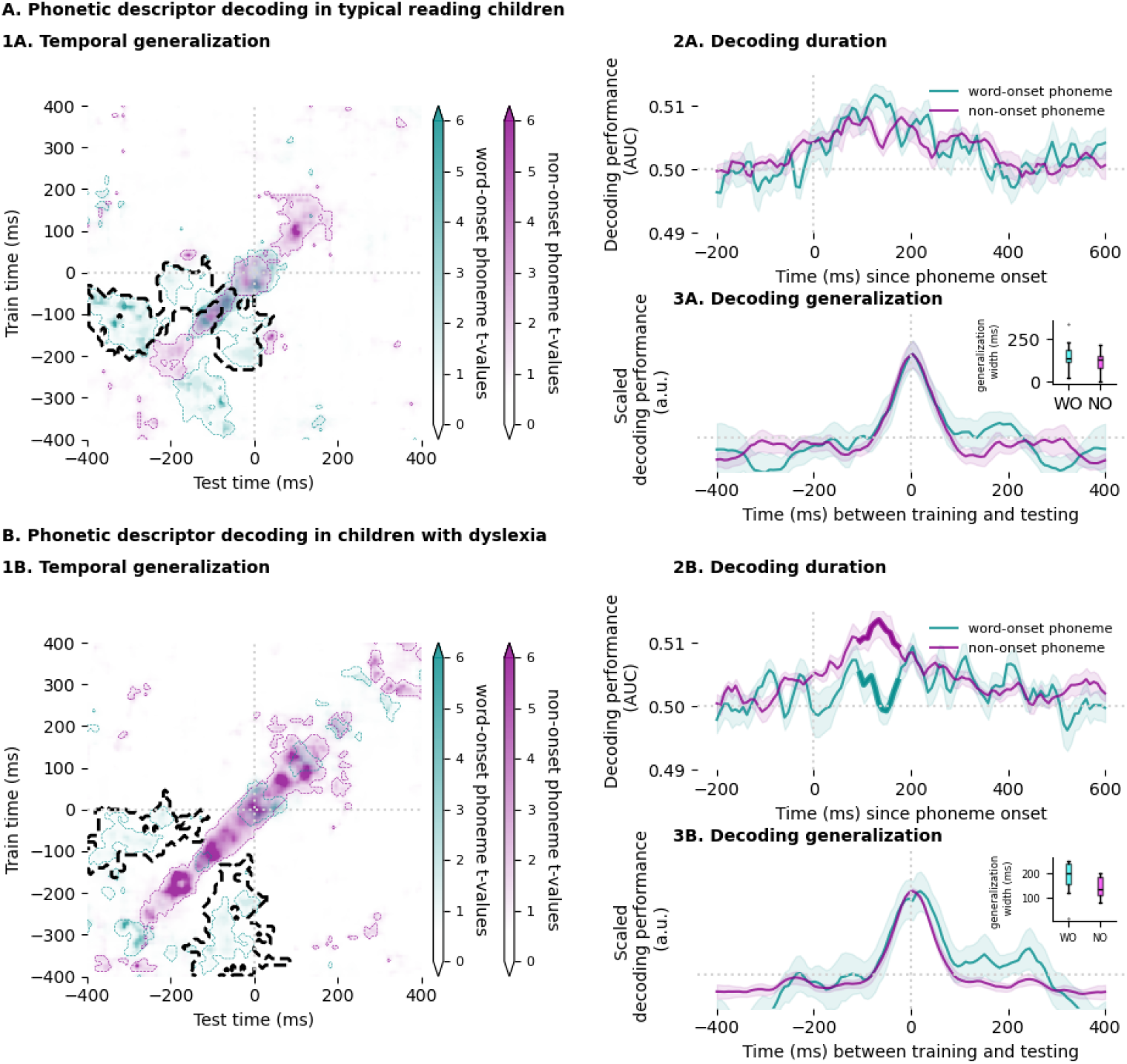
Neural dynamics of word-onset and non-onset phoneme processing in typical readers and children with dyslexia. **Panel 1A**: Temporal generalization of the average phonetic descriptor for word-onset (teal) and non-onset (magenta) phonemes in typical readers. Colored contours indicate clusters with decoding performance significantly above chance; the black, striped contour marks significant differences between conditions. **Panel 2A**: No significant difference in decoding duration between word-onset and non-onset phonemes in typical readers. **Panel 3A**: No significant difference in decoding generalization between conditions in typical readers. **Panel 1B**: Temporal generalization for word-onset and non-onset phonemes in children with dyslexia, with contours indicating significant decoding clusters. The black, striped contour highlights significant differences between conditions. **Panel 2B**: Decoding duration differed significantly between word-onset and non-onset phonemes in children with dyslexia; thicker line segments mark significant time points. **Panel 3B**: Decoding generalization also differed significantly in children with dyslexia, with thicker segments indicating significant intervals.

